# Acetylation is required for NLRP3 self-aggregation and full activation of the inflammasome

**DOI:** 10.1101/2019.12.31.891556

**Authors:** Kai Zhao, Yening Zhang, Xueming Xu, Liping Liu, Lingmin Huang, Ruiheng Luo, Jing Li, Ningjie Zhang, Ben Lu

**Affiliations:** Department of Hematology and Key Laboratory of Non-resolving Inflammation and Cancer of Hunan Province, the Third Xiangya Hospital, Central South University, Changsha, Hunan Province, 410000 P. R. China; Department of General Surgery, the Third Xiangya Hospital, Central South University, Changsha, Hunan Province, 410000 P. R. China; Department of Blood Transfusion, The Second Xiangya Hospital, Central South University, Changsha, Hunan Province, 410000 P. R. China; Key Laboratory of Medical Genetics, School of Biological Science and Technology, Central South University, Changsha, Hunan Province, 410000 P. R. China; Key Laboratory of sepsis and translational medicine, School of Basic Medical Science, Central South University, Changsha, Hunan Province, 410000 P. R. China; Department of Pathophysiology, School of Basic Medical Science, Jinan University, Guangzhou, Guangdong Province, 510632 P. R. China

**Keywords:** acetylation, NLRP3 inflammasome, assembly process, self-aggregation, KAT5

## Abstract

The full activation of NLRP3 inflammasome needs two sequential signals: the fist priming signal and the second assembly signal. Various stimuli including infections and stress signals can provide the assembly signal. However, how NLRP3 detects diverse stimuli and becomes fully activated remain largely unknown. In this study, we found the second signal specially triggers the acetylation of NLRP3, which facilitates the aggregation of NLRP3 and its interaction with ASC and NEK7, thus promoting the assembly of inflammasome. Meanwhile, by employing pharmacological and molecular approaches, we identified KAT5 as a regulator of NLRP3 acetylation and activation. Furthermore, KAT5 specific inhibitor-NU9056 exhibited a robust suppressive effect on NLRP3 inflammasome both *in vitro* and *in vivo*. Thus, our study reveals a new mechanism for NLRP3 full activation and suggests targeting NLRP3 acetylation may provide a new approach for treatment of NLRP3 associated diseases.

## Introduction

NLRP3, which belongs to NLR family, can detect a broad range of microbial motifs, endogenous danger signals and environmental irritants(*1–3*). Upon activation, NLRP3 forms an inflammasome complex with the adapter ASC, leading to the activation of caspase-1, release of proinflammatory cytokines, and cell death. Aberrant NLRP3 inflammasome is regard as initiator and promotor of series of diseases including inflammatory, metabolic, degenerative and aging-related diseases(*1–3*).

Currently, it is well accepted that NLRP3 inflammasome activation needs two sequential steps, namely priming and assembly(*3–5*). Pathogen-associated molecule patterns (PAMPs), such as LPS, provide the first signal, which promotes the expression of NLRP3 and IL-1β. Diverse stimuli, such as ATP or nigericin, provide the second signal, triggering the assembly of NLRP3 inflammasome. Although there are several NLRP3 inflammasome activation models have been proposed including potassium efflux, mitochondrial damage, ROS production, lysosomal disruption and Trans-Golgi disassembly to explain the common pathway for different stimuli(*3–5*), it still remains unclear that how these diverse stimuli result in the assembly of NLRP3 inflammasome. Further efforts are still needed to elucidate the precise mechanism by which the NLRP3 senses these numerous stimuli.

Recent studies have provided a molecular mechanism for NLRP1B activation(*6, 7*), in which they have found ubiquitination on N terminal of NLRP1B promotes the degradation of N terminus and frees the NLRP1B C terminus to activate caspase-1, suggesting modulation of post-translational modifications (PTMs) is critical for inflammasome activation. Although there are several PTMs on NLRP3 were reported, such as ubiquitylation, phosphorylation and sumoylation(*3, 8, 9*), these studies mostly focused on the PTMs induced by the priming signal (*10–12*), the second signal induced PTMs on NLRP3 and associated activities changed remains less defined.

Here, we discovered the second signal specially triggered the acetylation on Lys24 of NLRP3, which was indispensable for the assembly of NLRP3 inflammasome. It boosted the aggregation of NLRP3 and promoted associated interaction with ASC and NEK7. By employing pharmacological and molecular approaches, we identified KAT5 as a regulator of NLRP3 acetylation and activation. We also found NU9056, one specific KAT5 inhibitor could suppress the NLRP3 inflammasome activation both *in vitro* and *in vivo*. Thus, our study uncovers that acetylation boosts NLRP3 self-aggregation and full activation of the inflammasome in response to diverse stimuli during assembly process.

## Results

### Identification of NLRP3 acetylation

Since acetylation plays a pivotal role in various biological processes (*13, 14*), we asked whether it participates in NLRP3 inflammasome activation. We treated primary macrophages with LPS, LPS+ATP, and LPS+nigericin, then collected the whole cell lysis extracts to detect the acetylation level by using an antibody to all acetylated lysines, we found that the acetylation level obviously increased upon LPS+ATP or LPS+nigericin stimulation (Figure S1A), indicating that acetylation has a regulatory role in NLRP3 inflammasome activation. Intriguingly, we noticed that acetylation level changed significantly between 55Kd and 130Kd, in which only NLRP3 is within the molecular weight. Thus, we decided to figure out whether NLRP3 is acetylated. For this, we adopted a standard reconstitution system in HEK293T cells (*15*), and stimulated with nigericin. Since maturation of extracellular IL-1β is regarded as an indictator of NLRP3 activity (Figure S1B), this reconstitution system provided a relevant model to study NLRP3 inflammasome. After that, we adopted an acetylproteomic approach to identify the acetylation sites in NLRP3. Flag-tagged NLRP3 was immunoprecipitated and analyzed by liquid chromatography-mass spectrometry (Figure S1C and S1D). Three acetylation sites (Lys24, Lys234 and Lys875) were identified. Among them, Lys234 and Lys875 were the common sites in the two groups (non-treated and nigericin treated groups), while the Lys24 was exclusively identified in nigericin treated group.

To further determine the role of these acetylation sites in NLRP3 inflammasome activation, we replaced lysine at position 24, 234, or 875 with glutamine (K24Q, K234Q, or K875Q,) or arginine (K24R, K234R or K875R) to mimic the acetylated or nonacetylated state of the lysine residue, respectively. Then the function of each NLRP3 variant was examined in HEK293T cells reconstituted with NLRP3 inflammasome. The K24R and K24Q NLRP3 mutants were unable to promote IL-1β cleavage, while the others have the similar promoting effect as wild type NLRP3 (Figure 1A and 1B). The Lys24 residue, as the unique site identified in nigericin treated group, immediately has attracted our attention. The Lys24 residue is located in the pyrin domain (PYD) and conserved across many species (Figure 1C). To further address the functional importance of Lys24 in activation of NLRP3 inflammasome, we re-expressed of NLRP3-K24R or NLRP3 wild type (WT) in NLRP3^−/−^ iBMDMs by lentivirus, then stimulated with ATP or nigericin and found that re-expression of NLRP3-K24R impaired the IL-1β but not TNF-α secretion compared to re-expression of WT NLRP3 (Figure 1D). Collectively, these results suggested lys24 acetylation of NLRP3 was critical for its full activation.

**Figure 1.**
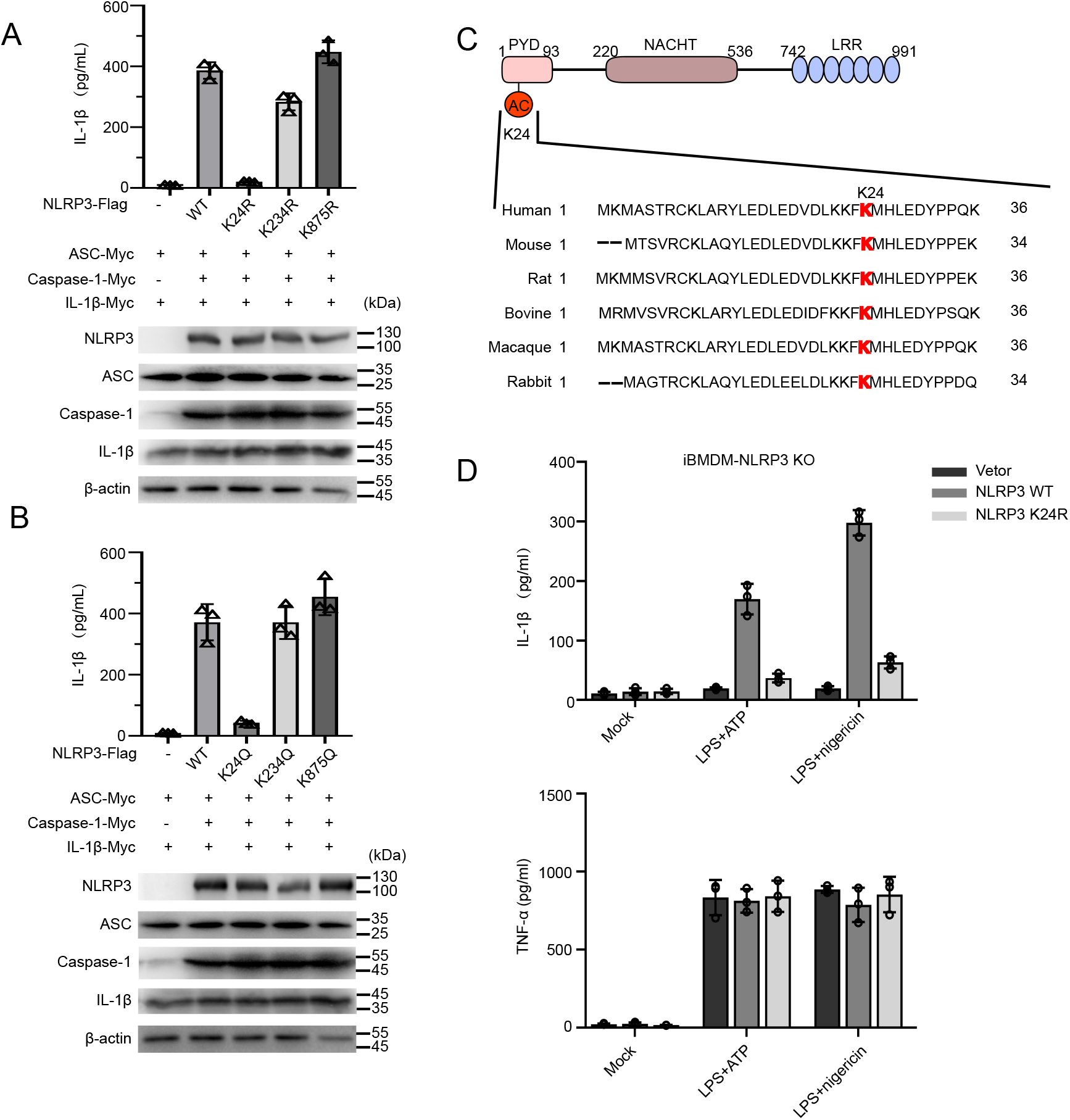
NLRP3 Acetylation at Lys24 is indispensable for NLRP3 inflammasome activation. (A and B) ELISA of IL-1β in supernatants and Immunoblot of whole cell lysis from HEK293T cells reconstituted with NLRP3 inflammasome and stimulated with nigericin. For NLRP3 inflammasome reconstitution, HEK293T cells were transiently transfected with Flag-tagged WT NLRP3 or its nonacetylation mimetic (K-to-R) mutations (A) or K-to-Q mutations (B) and Myc-tagged pro-caspase-1, pro-IL-1β, ASC plasmids. (C) Scheme of NLRP3 domain structure and the alignment of NLRP3 orthologs. The acetylated lysine residue is underlined in red. (D) NLRP3 Knockout iBMDMs transduced with lentivirus expressing WT or K24R NLRP3, then primed with LPS, and stimulated with ATP or nigericin. ELISA of IL-1β and TNF-α in supernatants were measured. Experiments were repeated at least three times, and representative data are shown. Values, mean± SD.

### NLRP3 is acetylated during the assembly process

Next, we studied when NLRP3 became acetylated during inflammasome activation. By treated primary macrophages with different NLRP3 inflammasome stimuli, including ATP (K^+^-efflux dependent), MSU (lysosome disruption dependent) and Imiquimod (K^+^-efflux independent) (*3*), we found NLRP3 was indeed acetylated following the second signal (Figure 2A), while an inapparent acetylation band was shown in LPS treated group, consistent with the results of mass spectrometry. In addition, activation of AIM2 or NLRC4 inflammasome did not induce NLRP3 acetylation (Figure 2B).

**Figure 2.**
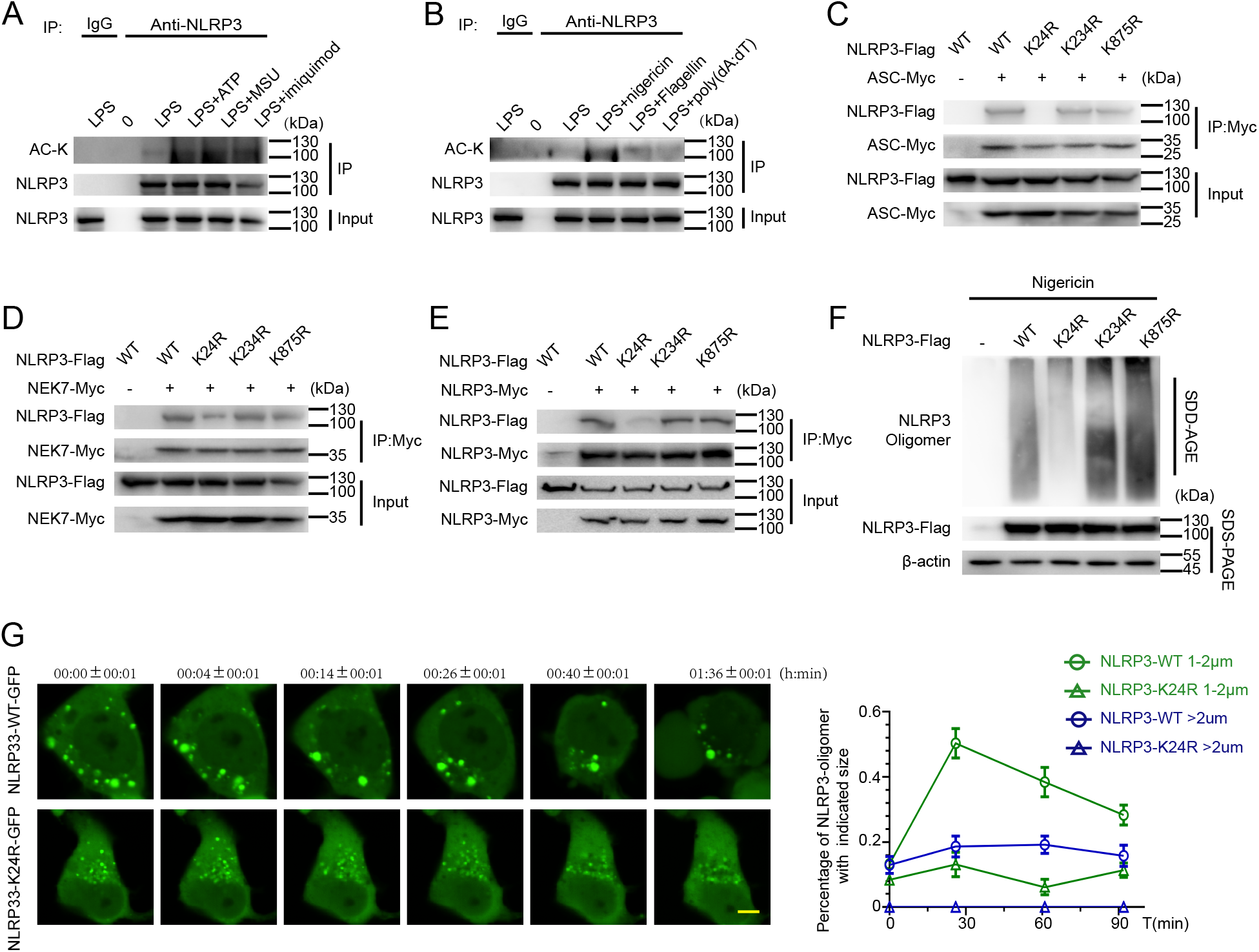
NLRP3 K24 acetylation promotes the inflammasome assembly. (A and B) Immunoblot analysis of acetylation level of NLRP3 immunoprecipitated from total lysis of peritoneal macrophage treated with kinds of NLRP3 inflammasome stimuli (A) or other inflammasome stimuli (B). (C-E) Immunoblot analysis of ASC(C), NEK7(D), or NLRP3(E) in HEK293T cells transfected to overexpress Myc-tagged ASC, NEK7, or NLRP3 alone or together with Flag-tagged NLRP3 or K24R NLRP3, assessed before (Input) or after (IP) immunoprecipitation with antibody to Myc. (F) Immunoblot analysis of NLRP3 oligomerization by SDD-AGE or SDS-PAGE assay in HEK293T cells transfected with WT or K24R NLRP3 plasmid and then stimulated with nigericin. (G) Microscopy imaging of NLRP3 oligomerization in HEK293T cells transfected with WT or K24R NLRP3-GFP plasmid and stimulated with nigericin. Shown are representative of time-lapse cell images taken from 5 min after nigericin addition (left). The percentage of NLRP3 oligomer was quantified (right). Scale bar, 5μm. For videos of two representative cells, see Supplementary Videos 1 and 2. Experiments were repeated at least three times, and representative data are shown. Values, mean± SD.

### Lys24 acetylation in NLRP3 regulates NLRP3 self-aggregation

We then investigated how NLRP3 Lys24 acetylation regulates NLRP3 inflammasome full activation. Lys24 is located in the PYD domain of NLRP3, previous studies have showed that PYD domain mediates the association between NLRP3 and ASC, which is essential for the assembly of the inflammasome(*3, 4*). We asked whether Lys24 acetylation of NLRP3 affects inflammasome assembly. We transfected those nonacetylated mutation NLRP3 plasmids and found the interrupted interaction between NLRP3-K24R and ASC compared with NLRP3-K234R, NLRP3-K875R, and NLRP3 WT (Figure 2C). Recent work have suggested NEK7 is an essential factor for NLRP3 inflammasome assembly(*16–18*), therefore, we detected the association between NLRP3 and NEK7and found that K24R mutation of NLRP3 also impaired their interaction (Figure 2D). Thus, these data suggested Lys24 in NLRP3 promote the assembly of inflammasome by affecting the association between NLRP3 and ASC or NEK7.

Since self-aggregation of NLRP3 occurred earlier than the association between NLRP3 and ASC during the inflammasome assembly(*3, 4*), we then speculated whether Lys24 acetylation regulates the NLRP3 aggregation. We firstly detected the interaction between Flag-NLRP3 and Myc-NLRP3 and found the association was decreased when K24R mutant was introduced (Figure 2E). Previous studies have showed that HEK293T-expressed NLRP3 could form oligomers in response to nigericin treatment(*10, 19*). Accordingly, we expressed WT NLRP3 and K24R NLRP3 in HEK293T cells, then treated them with nigericin and found the NLRP3 oligomerization was robustly decreased in HEK293T-expressed K24R NLRP3 by adopting semidenaturing detergent agarose gel electrophoresis (SDD-AGE) (Figure 2F), a method for detecting large protein oligomers in studying prions(*20, 21*). This was further confirmed by fluorescence microscopy. Oligomerization of NLRP3 was remarkably abrogated when K24R mutant was introduced in HEK293T cells stimulated with nigericin. (Figure 2G and supplementary video1 and 2). Taken together, these data demonstrated K24 acetylation in NLRP3 regulates NLRP3 inflammasome assembly by influencing NLRP3 self-aggregation.

### KAT5 mediates the acetylation of NLRP3

To explore which acetyltransferase was responsible for the acetylation of NLRP3, we screened the acetyltransferase by using a series of acetyltransferase inhibitors. To examine the effect of acetyltransferase inhibitor on the second signal of NLRP3 inflammasome We treated LPS primed macrophages with these inhibitors, then stimulated with nigericin. Intriguingly, we found NU9056 exhibited a robust inhibition of NLRP3 inflammasome with less IL-1β secretion but not TNF-α, while the others had no such obvious suppressive effect (Figure S2A).

Since NU9056 is an inhibitor of KAT5(*22, 23*), we next to study the role of KAT5 in NLRP3 acetylation. We firstly detected the interaction between KAT5 and NLRP3 in HEK293T cells, and confirmed the strong interaction between KAT5 with NLRP3 (Figure 3A), but not with ASC or caspase-1 (Figure S2B and S2C). This association was also verified in primary macrophages (Figure 3B). Meanwhile, we noticed that the association between NLRP3 and KAT5 was mostly increased upon nigericin treatment (Figure 3B), which was also supported by confocal microscopy assays (Figure 3C), indicating KAT5 regulates NLRP3 activity during the second signal process. Furthermore, we demonstrated KAT5 directly interacted with NLRP3 by applying a GST pull-down assay (Figure 3D). Then we examined the role of KAT5 in NLRP3 acetylation. We found the acetylation level of NLRP3 was obviously reduced in KAT5 knockdown or NU9056 treated macrophages (Figure 3E and Figure S2D). In vitro acetylation assay further demonstrated KAT5 acetylated NLRP3 (Figure 3F). Taken together, these data implied that KAT5 interacts with NLRP3 and directly mediates the acetylation of NLRP3.

**Figure 3.**
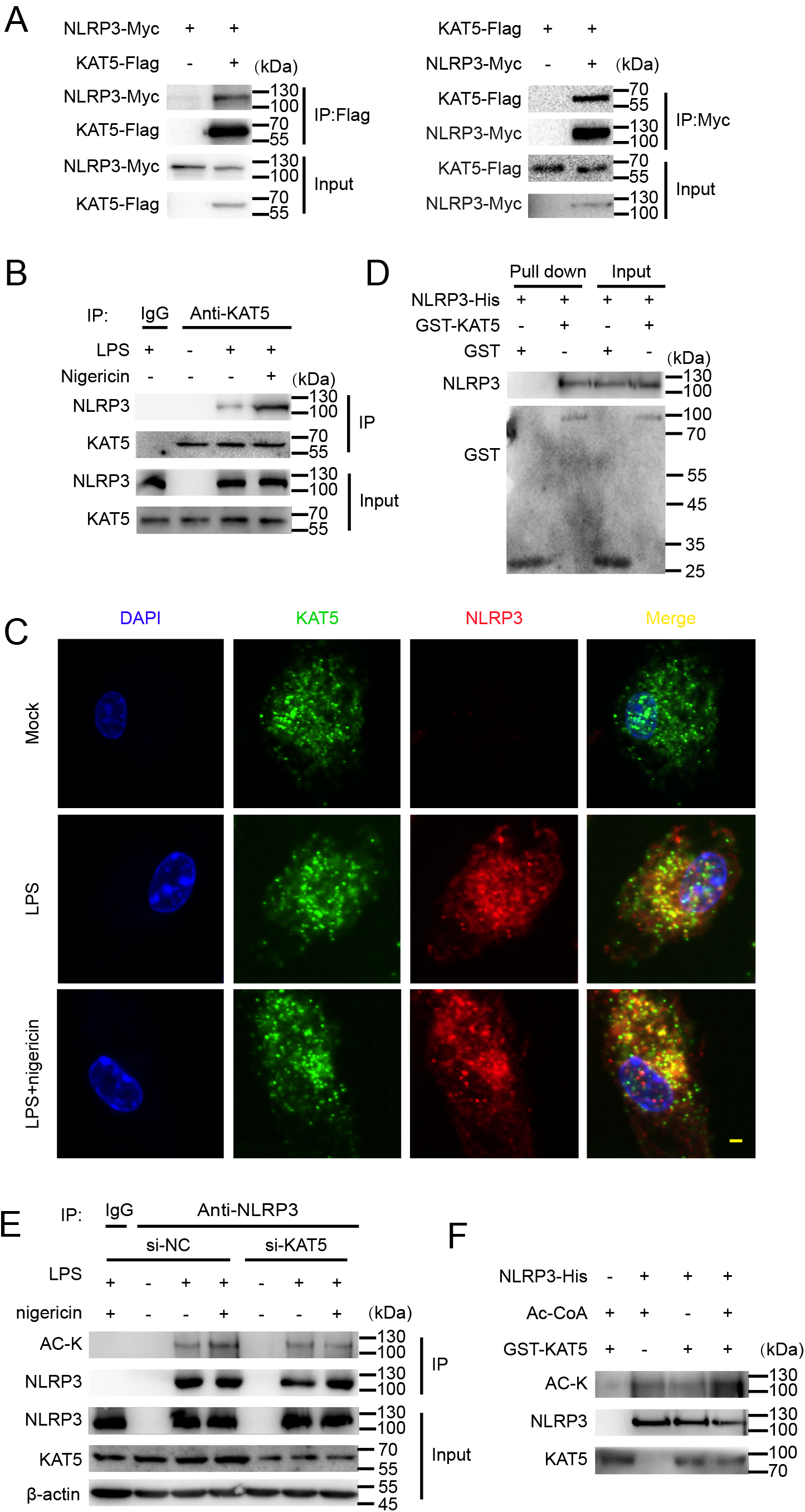
KAT5 mediates NLRP3 acetylation. (A) Immunoblot analysis of NLRP3 or KAT5 in HEK293T cells transfected to overexpress Myc-tagged NLRP3 alone or with Flag-tagged KAT5, assessed before (Input) or after (IP) immunoprecipitation with antibody to Myc or Flag. (B) Immunoblot analysis of the endogenous NLRP3 or KAT5 in peritoneal macrophages treated with indicated stimuli. assessed before (Input) or after immunoprecipitation (IP) with IgG or antibody to KAT5. (C) Representative images taken from confocal microscopy of intracellular Co-localization of KAT5 and NLRP3 in peritoneal macrophages upon indicated stimuli. Scale bar, 2μm. (D) GST-Pull down analysis of the interaction between GST-KAT5 and His-NLRP3. (E) Immunoblot analysis of acetylation level of NLRP3 immunoprecipitated from total lysis of peritoneal macrophages with indicated stimuli after transfected with NC-siRNA or KAT5-siRNA. (F) Immunoblot analysis of acetylation level of NLRP3 *in vitro*. The recombinant His-NLRP3 was incubated with GST-KAT5 in the presence or absence of acetyl-CoA (Ac-CoA). Experiments were repeated at least three times, and representative data are shown. Values, mean± SD.

### KAT5 promotes NLRP3 inflammasome activation

To further examine the role of KAT5 in NLRP3 inflammasome activation, we silenced KAT5 in primary macrophages and found that knockdown of KAT5 remarkably suppressed NLRP3 inflammasome activation, reflected by IL-1β secretion, caspase-1 maturation and cell death (Figure 4A–4C), while the activation of AIM2 and NLRC4 inflammasomes were not affected (Figure 4A and 4B). Since KAT5 could regulate gene expression in a transcriptional manner, we found knockdown of KAT5 had no effect on the mRNA expression of NLRP3 inflammasome components (Figure S3A–S3D). Similarly, knockdown of KAT5 in iBMDMs by short haipin RNAs (shRNAs) resulted in impaired NLRP3 inflammasome activation but not AIM2 or NLRC4 inflammasomes (Figure S4A–S4C). ASC nucleation-induced oligomerization and ASC-speck formation were also inhibited by KAT5 knockdown (Figure 4D and 4E). Then we examined whether KAT5 affects the NLRP3 inflammasome assembly. We found the association between NLRP3 and ASC or NEK7 was markedly disrupted in the absence of KAT5 (Figure 4F–4G and Figure S4D–S4E), and the endogenous NLRP3 oligomerization was also robustly decreased upon KAT5 knockdown (Figure 4H and Figure S4F). Meanwhile, NU9056 treatment got the similar suppressive effect on NLRP3 oligomerization (Figure S4G). Thus, these data demonstrated that KAT5 promotes NLRP3 inflammasome activation via regulating the self-aggregation of NLRP3 and its interaction with ASC and NEK7.

**Figure 4.**
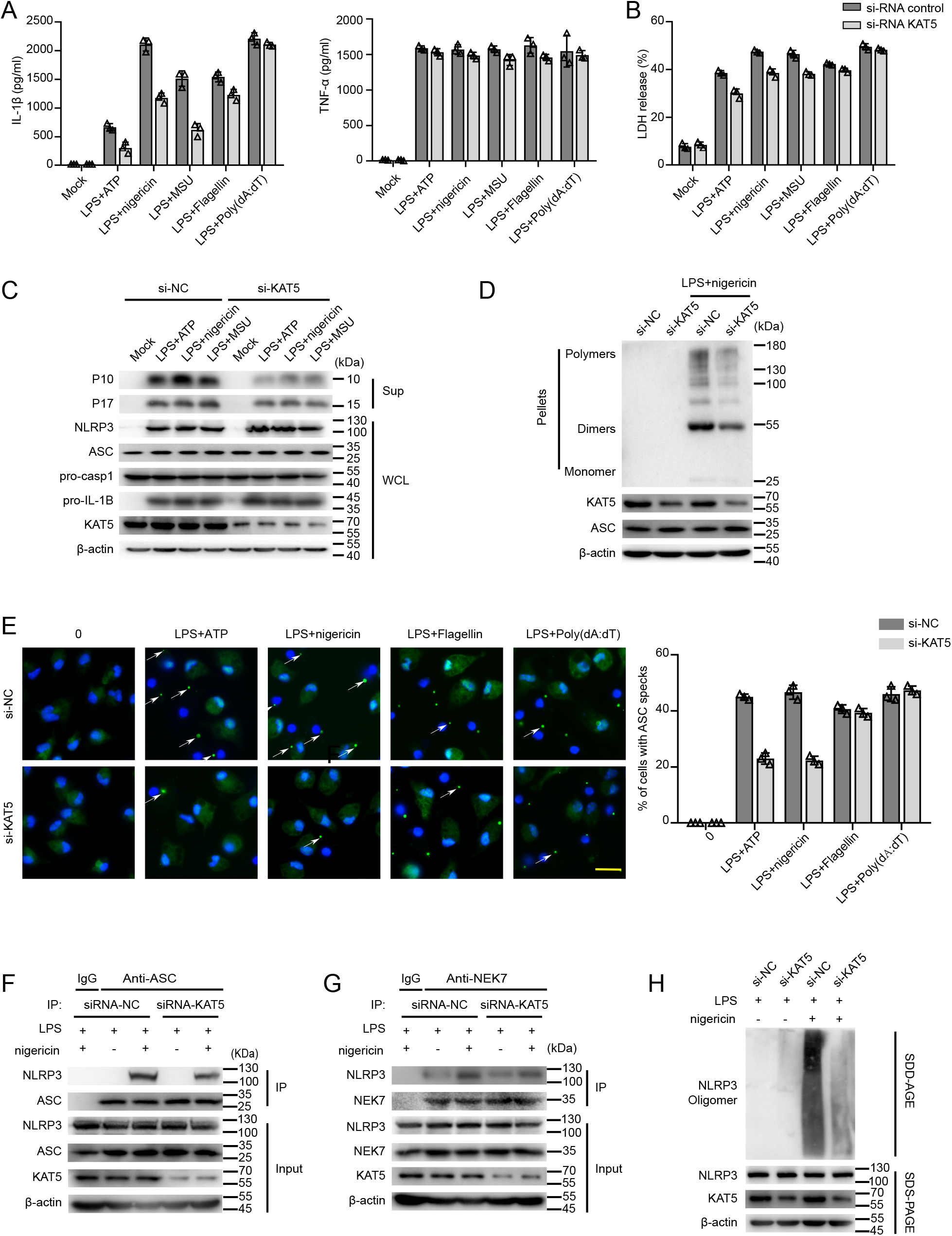
KAT5 specifically promotes NLRP3 inflammasome activation. (A-C) Effects of KAT5 in NLRP3 inflammasome activation in peritoneal macrophages. ELISA of IL-1β and TNF-α (A), release of LDH (B) in supernatants and immunoblot analysis of extracts (C) from peritoneal macrophages with indicated stimuli after transfected with NC-siRNA or KAT5 siRNA. (D and E) Effects of KAT5 in NLRP3-mediated ASC oligomerization (D)and speck formation (E) in peritoneal macrophages with indicated stimuli after transfected with control siRNA or KAT5 siRNA. Representative images of ASC speck subcellular distribution and quantification of ASC speck formation by the number of cells with ASC specks relative to the total number of cells from 5 random fields each containing 50 cells on indicated stimulation. Scale bar, 20 μm. (F and G) Effects of KAT5 on the interaction between NLRP3 and ASC or NEK7. Primary macrophages treated with indicated stimuli after transfected with NC or KAT5-siRNA, immunoprecipitation was performed to detect the interaction between NLRP3 and ASC(F) or NEK7(G). (H) Effects of KAT5 on NLRP3 oligomerization. Immunoblot analysis of NLRP3 oligomerization by SDD-AGE or SDS-PAGE assay in peritoneal macrophage treated with indicated stimuli after transfected with NC-siRNA or KAT5 siRNA. Experiments were repeated at least three times, and representative data are shown. Values, mean±SD.

### Acetylation inhibitor-NU9056 suppresses NLRP3 inflammasome in vivo

As acetylation plays a crucial role in NLRP3 inflammasome activation, we next to determine whether inhibition of NLRP3 acetylation could alleviate NLRP3 inflammasome related inflammatory responses *in vivo*. Two NLRP3-dependent models were adopted-LPS induced acute systemic inflammation model MSU induced peritonitis model (*10, 24*). We pre-treated mice with KAT5 inhibitor-NU9056 before subjected to the two models. In the systemic inflammation model, mice pre-treated with NU9056, exhibited markedly reduced IL-1β secretion but little change in TNF-α or IL-6 production, compared to the saline pre-treated group (Figure 5A–5C), and their lungs also exhibited less severe tissue damage and diffuse inflammation (Figure 5D). Mice pre-treated with NU9056 were also refractory to MSU-induced peritonitis, and exhibited reduced peritoneal exudate cells relative to saline pre-treated mice (Figure 5E and 5F). The suppressive effect of NU9056 is dependent on NLRP3 inflammasome, as it had no effect on AIM2 or NLRC4 inflammasome (Figure S5A–S5C). Collectively, these findings indicated that targeting NLRP3 acetylation might provide a new approach for treatment of NLRP3 inflammasome associated inflammatory diseases and NU9056 might be a candidate small molecular.

**Figure 5.**
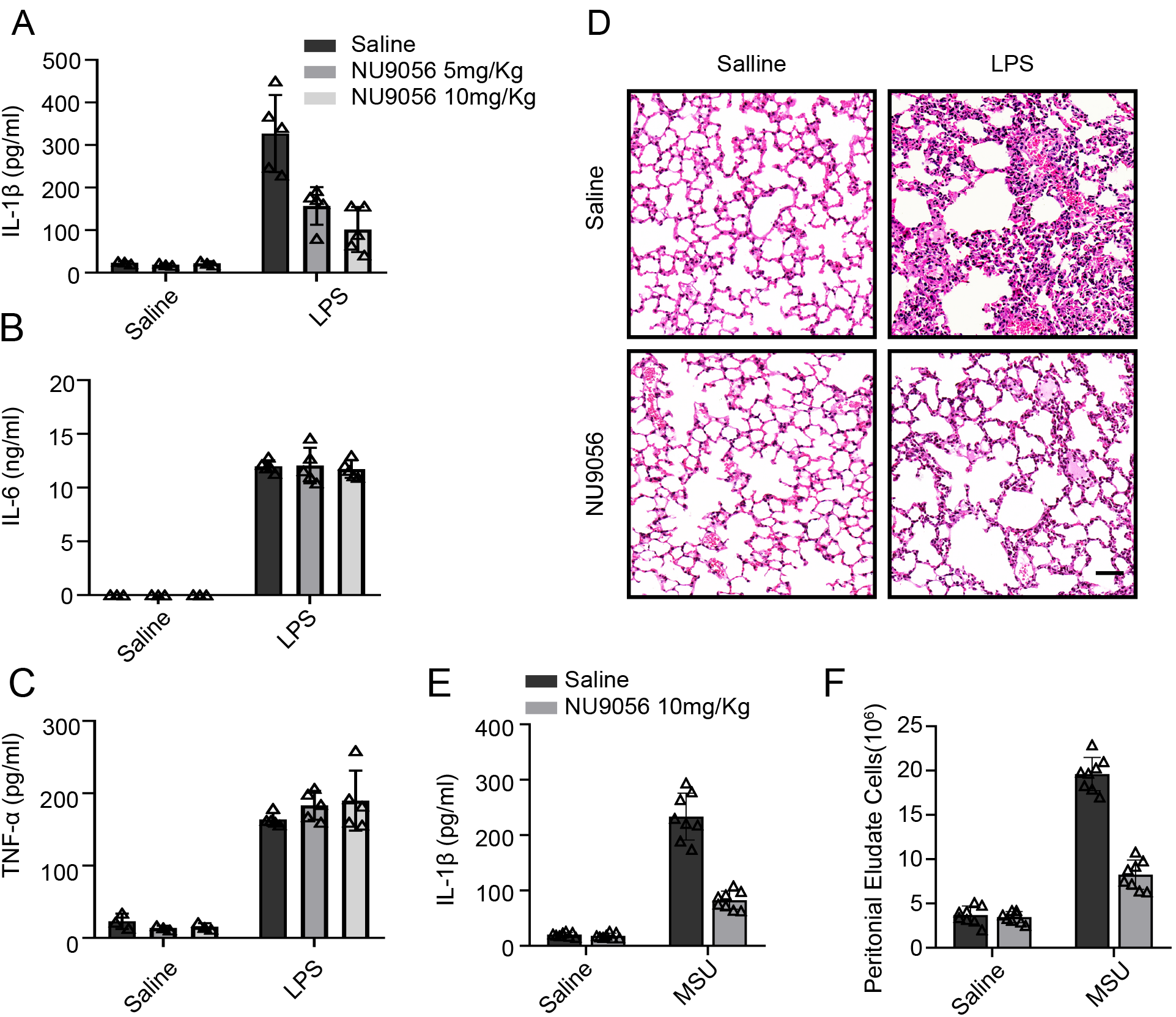
Acetylation inhibitor-NU9056 suppresses NLRP3 inflammasome *in vivo*. (A-D) Effects of NU9056 in LPS induced systemic inflammation. Wild-type C57BL/6 mice were intraperitoneal (i.p.) injection of Saline or NU9056 (5 or 10 mg/kg) 30min followed by i.p. injection of LPS (20mg/kg) for 8h. ELISA analysis of serum IL-1β, IL-6(B), TNF-α (C) and representative H&E images of lung sections (D) Scale bar, 50μm. (E and F) Effects of NU9056 in MSU-induced peritonitis. Wild-type C57BL/6 mice were intraperitoneal (i.p.) injection of Saline or NU9056 (5 or 10 mg/kg) 30min followed by i.p. injection of MSU (1mg) for 6h. ELISA analysis of IL-1β (E) and quantification of peritoneal exudate cells (F) in the peritoneal cavity fluid. Experiments were repeated at least three times, and representative data are shown. Values, mean±SD.

## Discussion

By employing acetylproteomic approach, our study identified an acetylation site-Lys24 of NLRP3, triggered by the second signal, is indispensable for the NLRP3 inflammasome assembly through regulating the self-aggregation of NLRP3 and associated interaction with ASC and NEK7. By screening the acetyltransferase, we identified KAT5 as a regulator of NLRP3 acetylation and activation. Furthermore, KAT5 inhibitor-NU9056 exhibited a robust inhibitory effect on NLRP3 inflammasome both *in vitro* and *in vivo*, providing a therapeutic potential for NLRP3 associated diseases.

NLRP3 inflammasome has attracted most attention among the numerous inflammasomes, however, the underlying mechanism of NLRP3 detecting diverse stimuli still remains unknown(*1–3*). Although previous studies have established several models, including potassium efflux, mitochondrial damage, ROS production, lysosomal disruption and Trans-Golgi disassembly, to explain the common pathway for different stimuli(*3–5*), some of which are interrelated and overlapping, and the data are sometimes conflicting(*3*). Recent works have suggested PTMs play a pivotal role in molecular mechanism of NLRP1b activation(*6, 7*), making us pay more attention to the regulatory PTMs of NLRP3. Ubiquitination is the first PTMs reported in NLRP3 regulation, the authors have shown that BRCC3 mediates the deubiquitination of NLRP3 upon stimuli and contributes to its full activity(*25*). Since then, a number of E3 ligases have been identified for promoting K48-linked ubiquitination and degradation of NLRP3, such as PAI2(*26*), MARCH7(*24*), Trim31(*27*) and FBXL2(*28*). Except for ubiquitination, phosphorylation of NLRP3 by different kinases have been widely studied(*3, 8, 9*). Undoubtedly, these studies broadening our understanding of the NLRP3 activation, however, most of which focused on how priming signal-induced phosphorylation of NLRP3 and associated activity (*10–12*). In our study, we focused on the second signal-induced PTMs during NLRP3 activation. We found the second signal trigger the acetylation of NLRP3, and also identified the acetylation site Lys24 of NLRP3. When we mutated the K24 to R24, the self-aggregation of NLRP3 was markedly inhibited and the associated interaction with ASC and NEK7 was also significantly interrupted, suggesting acetylation is indispensable for NLRP3 inflammasome assembly and full activation. Recent work suggested the second signal triggers desumolation of NLRP3(*29*) lead to the NLRP3-dependent ASC oligomerization, caspase-1 activation and IL-1β release, consistent with our study that PTMs occurred at the second signal contribute to the assembly of NLRP3 inflammasome. Thus, our study suggests diverse stimuli fully activates NLRP3 inflammasome by triggering the acetylation of NLRP3. However, whether exists other PTMs induced by the second signal needs further investigation.

Lys24, the acetylation site identified in our study, is located in the PYD domain. Recent work has showed that phosphorylation of Ser3 in the PYD domain of NLRP3 disturb the interaction between NLRP3 and ASC by introducing negative charge(*11*). Consistently, acetylation of the lysine side chain can neutralize the positive charge, we speculated that acetylation of Lys24 might also affect charge-charge interaction for NLRP3 aggregation. Interestingly, Lys24 is conserved across many species (in human is lys26), but there is no related report on this site mutation in human diseases, owing to the loss-of-function mutation is less common than the gain-of-function mutation in NLRP3 related diseases, which still needs to be further investigated.

KAT5 (also known as Tip60), belongs to MYST family of histone acetyltransferase, plays a pivotal role in diverse cellular activities including chromatin remodeling, DNA repair, gene transcription, apoptosis, and tumorigenesis(*30, 31*). Previous studies have suggested that KAT5 contributes to chronic inflammatory responses(*32, 33*), such as rheumatoid arthritis (RA) and allergy, by regulating the Foxp3 expression in regulatory T cells and STAT6 expression in B cells, respectively (*32–34*). However, the role of KAT5 in acute inflammatory responses mediated by innate immune cells is largely unknown. In our study, we revealed that KAT5 could induce the acetylation of NLRP3 and promote its full activity, contributing to NLRP3 associated inflammatory responses. As NLRP3 is a non-histone protein, our finding also supports previous study that KAT5 could mediate non-histone proteins acetylation (*30, 31, 35*). Thus, we have uncovered a novel role of KAT5 in innate immune responses.

Together with others findings that targeting the PTMs of NLRP3 may providing a potential for treating NLRP3 associated diseases(*8, 9, 12, 25*), In our study, KAT5 specific inhibitor-NU9056 could suppress the NLRP3 inflammasome activation both *in vitro* and *in vivo* by inhibiting NLRP3 acetylation. Taken together, our study adds a new view for NLRP3 full activation and suggests targeting NLRP3 acetylation may provide a new approach for treatment of NLRP3 associated diseases.

## Acknowledgments

We thank Dr. Feng Shao (National Institute of Biological Sciences) for providing immortalized mouse macrophages. We thank Qianqian Xue for assisting in raising the animals. This work was supported by National Natural Science Foundation of China (81801967), Innovation-riven Project of Central South University (2018CX030,2019CX013).

## Author Contributions

K.Z., Y. Z., X.X., L.L., L.H., R.L., J.L. and N.Z., performed the experiments; K.Z. and Y. Z. analyzed the results; K.Z. and B. L. designed the research; K.Z., Y. Z. and B. L. wrote the manuscript. K.Z. and B. L. supervised the whole project

## Materials and Methods

### Materials

The sources of the reagents and other materials used are listed in Supplementary Table1.

### Mice

Wild-type C57BL/6 mice (8-10 weeks old) were purchased from Hunan SJA Laboratory Animal Co. Ltd (Changsha, China). All animals were kept under SPF conditions. Animals experiments were conducted in accordance with the Institutional Animal Care and Use Committee of Central South University.

### Cell culture

Primary peritoneal macrophages from C57BL/6 mice were maintained in 1640 supplemented with 10% FBS. HEK293T cells were obtained from American Type Culture Collection (Manassas, VA), and iBMDMS were a kind gift from Dr. Feng Shao. The cells were cultured in Dulbecco’s modified Eagle’s medium (DMEM) supplemented with 10% 10% FBS. All the medium contains 100 U/ml penicillin and 100 μg/ml streptomycin.

### Inflammasome stimulation

Primary peritoneal macrophages and iBMDMs were seeded in 24-well (3×10^5^) or 6-well (1×10^6^) culture plates. Macrophages were primed with LPS (100 ng/ml) for 3 h followed by stimulation as follows: 5 mM ATP for1 hr, 10 ng/mL nigericin for 1 hr, 200 μg/mL MSU for 6 hr, 20μM Imiquimod for 1hr, 2 μg/mL Flagellin was transfected into macrophages by Lipofectamine 2000 for 1hr. 1 μg/mL Poly (dA:dT) using Lipofectamine 2000. iBMDMs primed with LPS (1 μg/mL) for 6 h followed by stimulation as follows: 5 mM ATP for1 hr, 10 ng/mL nigericin for 1 hr, 200 μg/mL MSU for 6 hr, 2 μg/mL Flagellin was transfected into macrophages by Lipofectamine 2000 for 1hr. 1 μg/mL Poly (dA:dT) using Lipofectamine 2000.

### Reconstitution of NLRP3 inflammasome in HEK293T cells

A standard reconstitution system in HEK293T cells was referred (*15*), in brief, HEK293T cells were seeded into 24-well plates at density of 2×10^5^ cells per well in complete cell culture medium overnight before transfection. The cells were transfected with plasmids expressing pro-IL-1β-Myc (200 ng/well), pro-caspase-1-Myc (100 ng/well), ASC-Myc (20 ng/well), NLRP3-Flag (200 ng/well) using Linear Polyethylenimine. 24-36 hours later, replace the medium with 250 μl DMEM cell culture medium, then10 μg/mL nigericin was added one hour before supernatant collection. Cells lysates were collected using cell lysis buffer (CST) added with EDTA-free protease inhibitor cocktail. The IL-1β in the supernatant was assessed by ELISA.

### Identification of NLRP3 acetylation by Mass spectrometry

HEK293T cells were transfected with Flag-tagged NLRP3 treated with nigericin or not. Then the cells were lysed in IP buffer (1% (v/v) Nonidet P-40, 50 mM Tris-HCl (pH 7.4), 50 mM EDTA, 150 mM NaCl) contained with a protease inhibitor cocktail. Cell extracts were centrifuged at 12,000 × g for 10 min at 4°C and the supernatants were immunoprecipitated with anti-Flag M2 beads (Sigma), and then dissolved in SDS loading buffer. Samples were denatured at 95°C for 10 min before SDS-PAGE. NLRP3 in gel was digested with trypsin, and subjected to LC/tandem MS (MS/MS) analysis in Jingjie PTM Biolab Co. Ltd. (Hangzhou,China). Briefly, the peptides were analyzed using Q Exactive^TM^ Plus mass spectrometry (Thermo) coupled to an ekspert EASY-nLC 1000 (Thermo). The resulting MS/MS data were processed using Proteome Discoverer 1.3. Tandem mass spectra were searched against the protein sequence of NLRP3. Mass error was set to 10 ppm for precursor ions and 0.02 Da for fragment ions. Carbamidomethyl on Cys were specified as fixed modification and oxidation on Met and acetylation were specified as variable modifications. Peptide confidence was set at high, and peptide ion score was set > 20.

### Plasmids and transfection

NLRP3, caspase-1, pro-IL-1β, ACS, and KAT5 full-length sequences were obtained from mouse peritoneal macrophage cDNA, then cloned into pcDNA3.1 vector that contained different tags. Deleted, truncated, and point mutants were generated by PCR-based amplification and the construct encoding the wild-type protein as the template. All constructs were confirmed by DNA sequencing. The primers were as follows: NLRP3 forward, 5’-AACGGGCCCTCTAGACTCGAGATGACGAGTGTCCGTTGCAAGCTGGCTCA GTA-3’, NLRP3 reverse, 5’-TAGTCCAGTGTGGTGGAATTCCCAGGAAATCTCGAAGACTATAGTCAGCTCA-3’, caspase-1 forward, 5’-AACGGGCCCTCTAGACTCGAGATGGCTGACAAGATCCTGAGGGCAAAGA-3’, caspase-1 reverse, 5’-TAGTCCAGTGTGGTGGAATTCATGTCCCGGGAAGAGGTAGAAACGTTTTG-3’, pro-IL-1β forward, 5’-AACGGGCCCTCTAGACTCGAGATGGCAACTGTTCCTGAACTCAACTGTGA-3’, pro-IL-1β reverse, 5’-TAGTCCAGTGTGGTGGAATTCGGAAGACACGGATTCCATGGTGAAGTCAA-3’, ASC forward, 5’-AACGGGCCCTCTAGACTCGAGATGGGGCGGGCACGAGATGCCATCCTGGA-3’, ASC reverse, 5’-TAGTCCAGTGTGGTGGAATTCGCTCTGCTCCAGGTCCATCACCAAGTAGG-3’, KAT5 forward, 5’-AACGGGCCCTCTAGACTCGAGATGGCGGAGGTGGGGGAGATAATCGAGGGCT-3’, KAT5 reverse, 5’-TAGTCCAGTGTGGTGGAATTCCCACTTTCCTCTCTTGCTCCAGTCTTTGGGA-3’. Plasmids were transiently transfected into HEK293T cells with Linear Polyethylenimine.

### NLRP3 reconstitution in NLRP3^−/−^ iBMDM cells

For reconstitution, NLRP3−/− iBMDM cells were transduced with virus stocks containing either a wild-type or K24R mutant NLRP3-encoding lentivirus. Virus was produced in HEK293T cells by co-transfection with pCDH-MCS-EF1-copGFP-NLRP3(wild type or mutants), pSPAX2, and pVSV-G with a ratio of 3:2:1. Virus-containing supernatants were filtered through a 0.45-μm-pore-size filter (Millipore) and supplemented with polybrene (8 μg/ ml) before adding to cells. GFP-positive cells were then sorted by flow cytometry (FACS Aria II, BD Biosciences).

### ASC Oligomerization

Peritoneal macrophages were stimulated with 100 ng/mL LPS for 3 hr, followed by stimulation with or without 10 ng/mL nigericin (1 hr). Then the cells were lysed with Triton Buffer [50 mM Tris-HCl (pH 7.5), 150 mM NaCl, 0.5% Triton X-100, 0.1 mM PMSF and EDTA-free protease inhibitor cocktail] for 10 min at 4°C. The lysates were centrifuged at 6,000 × g for 15 min at 4°C, the supernatant was collected and pellets were washed twice and re-suspended in 200 μL Triton Buffer, then added 2 mM disuccinimidyl suberate (DSS) and cross-linked for 30 min at 37°C. Samples were centrifuged and the pellets were dissolved in SDS loading buffer for western blotting.

### ASC Speck Formation

Peritoneal macrophages were seeded on chamber slides and allowed to attach overnight. The following day, the cells were treated with indicated stimulators. Then the cells were fixed in 4% paraformaldehyde followed by ASC and DAPI staining. Cells were visualized by fluorescence microscope (Nikon Ti2-U).

### CRISPR/Cas9-mediated generation of NLRP3^−/−^ iBMDM cells

CRISPR/Cas9 genomic editing for gene deletion according to the previous publication (*36*). Guide RNA sequences targeting NLRP3(5’-gtcctcctggcataccatag-3’) with BsmbI sticky end were annealed and inserted into the lentiviral vectors pLenti-CRISPR v2(Addgene #52961) digested with BsmBI (NEB). Lentivirus was produced in HEK293T cells by co-transfection of plenti-sgRNA plasmid, pSPAX2, and pMD2.G with a ratio of 3:2:1. Virus-containing supernatants were filtered through a 0.45-μm-pore-size filter (Millipore) and supplemented with polybrene (8 μg/ ml) before adding to cells. iBMDMs then were transduced with lentivirus encoding NLRP3 guide RNA. Stable transduced cells were selected with 5 mg/ml puromycin for 72 h, and single colonies were obtained by serial dilution and amplification. Clones were confirmed by DNA sequencing and also by immunoblotting with anti-NLRP3 antibodies.

### Knock down of KAT5 in macrophages and iBMDMs

For peritoneal macrophages, siRNA was transfected using Lipofectamine RNAiMAX (Thermo Fisher Scientific) according to the manufacturer’s instructions. The siRNA sequences for mouse KAT5 (5’-CCACACUGCAGUAUCUCAATT-3’), and the negative control (5’-UUCUCCGAACGUGUCACGUTT-3’) were chemically synthesized by Sangon Biotech Co., Shanghai, China

For iBMDMs, shRNA targeting KAT5 were from Genechem Co., Shanghai, China, the sequences were as follows: shKAT5-1(5’-CTGCAACGCCACTTGACCAAA-3’), shKAT5-2(5’-CTGCTTATTGAGTTCAGCTAT-3’), negative control(5’-TTCTCCGAACGTGTCACGT-3’). iBMDMs were transduced with lentivirus-KAT5-RNAi or ctrl virus. 48 hr later, the cells were selected by culture with 5 mg/mL puromycin. Single colonies were obtained by serial dilution and amplification.

### Immunofluorescence

Primary macrophages were stimulated with LPS for indicated hours, then cells were fixed with 4% paraformaldehyde for 15 min and permeabilized with 0.1% Triton X100 for 10 min. After blocking with 3% BSA for 1hr, cells were incubated overnight with anti-NLRP3 antibody (1:100 in PBS containing 3% BSA) and KAT5 antibody (1:100 in PBS containing 3% BSA), followed by staining with DyLight 488-labeled secondary antibody (Invitrogen) and Alexa Fluor 594-conjugated secondary Ab (Invitrogen) (1:50 in PBS containing 3% BSA). Nuclei were co-stained with DAPI (Invitrogen). Stained cells were viewed using a confocal fluorescence microscope (SpinSR10; Olympus).

### Quantitative PCR

Total RNA was extracted by using RNA Fast 200 kit according to the manufacturer’s instructions (FASTAGEN). Complementary DNA was synthesized by using TransScript All-in-One First-Strand cDNA Synthesis SuperMix for qPCR (TransGen Biotech) according to the manufacturer’s protocols. Quantitative PCR was performed using SYBR Green (Vazyme Biotech) on a LightCycler 480 (Roche Diagnostics), and data were normalized to β-actin expression. The 2^-ΔΔCT^ method was used to calculate relative expression changes. Gene-specific primers were as follows: KAT5 forward, 5’-TCCCGGTCCAGATCACACTC-3’;KAT5 reverse, 5’-ACCTTCCGTTTCGTTGAGCG-3’; NLRP3 forward, 5’-TGGATGGGTTTGCTGGGAT-3’, NLRP3 reverse, 5’-CTGCGTGTAGCGACTGTTGAG-3’; IL-1β forward, 5’-GCAACTGTTCCTGAACTCAACT-3’ IL-1β reverse, 5’-ATCTTTTGGGGTCCGTCAACT-3’; caspasse-1 forward, 5’-ACAAGGCACGGG ACCTATG-3’; reverse, 5’-TCCCAGTCAGTCCTGGAAATG-3’.ASC forward, 5’-CTTGTCAGGGGATGAACTCAAAATT-3’; reverse, 5’-GCCATACGACTCCAGATAGTAGC-3’. β-actin forward, 5’-AGTGTGACGTTGACATCCGT-3’; β-actin reverse, 5’-GCAGCTCAGTAACAGTCCGC-3’

### Immunoprecipitation and Western blot

HEK293T cells after transfection or peritoneal macrophages after stimulation, then lysed in IP buffer, added with a protease inhibitor cocktail, pre-cleared cell lysates were then subjected to anti-Flag M2 affinity gel or anti-Myc affinity gel for 2 hr or with specific Abs and protein G plus-agarose overnight, then washed four times with IP buffer. Immunoprecipitates were eluted by boiling with 1% (w/v) SDS sample buffer. For immunoblot analysis, cells were lysed with CLB buffer (CST) supplemented with protease inhibitor cocktail and PMSF, and then protein concentrations in the extracts were measured with a bicinchoninic acid assay (Pierce). Equal amounts of extracts were separated by SDS-PAGE, and then they were transferred onto nitrocellulose membranes for immunoblot analysis.

### GST Pulldown assay

Recombinant murine NLRP3-GST and KAT5-his proteins were purchased from Sino Biological Inc (Beijing, China), which were expressed by eukaryotic expression system. Incubate 4μg KAT5-His protein on a rotator with glutathione resin-immobilized GST or GST-NLRP3 in GST pull down buffer(20 mM Tris-HCl pH7.9, 150 mM NaCl, 0.5 mM EDTA, 10% glycerol, 1 mM DTT, 0.1% Triton X-100) for 2 hr at 4°C. Beads were washed 3 times with pull-down buffer before analyzed by immunoblot.

### In vitro acetylation assay

The acetylation assay was performed by incubating recombinant murine NLRP3-GST with or without KAT5-his protein in 40 μL of reaction buffer containing 50mM Tris-HCl (pH 9.0), 2mM MgCl2, 50mM NaCl, 0.5mM DTT and 0.2mM PMSF with or without 1.5mM NAD+ according to the previous research(*35*). After incubation for 1 hr at 30 °C, the reaction was stopped by addition of 10 μl of 5× SDS sample buffer. Samples were then subjected to SDS–PAGE and western blotting.

### SDD-AGE

The oligomerization of NLRP3 was analyzed according to the previous report (*21*). Cells were lysed with Triton X-100 lysis buffer (0.5% Triton X-100, 50 mM Tris-Hcl, 150 mM NaCl, 10% glycerol, 1 mM PMSF, and protease inhibitor cocktail), which were then resuspended in 1× sample buffer (0.5× TBE, 10% glycerol, 2% SDS, and 0.0025% bromophenol blue) and loaded onto a vertical 1.5% agarose gel. After electrophoresis in the running buffer (1× TBE and 0.1% SDS) for 1 h with a constant voltage of 80 V at 4°C, the proteins were transferred to Immobilon membrane (Millipore)for immunoblotting. 1× TBE buffer contains 89 mM Tris, pH 8.3, 89 mM boric acid, and 2 mM EDTA.

### ELISA assay for cytokines

Levels of IL-1β, TNF-α and IL-6 collected from cell culture and sera were determined using quantitative ELISA kits (eBioscience) according to the manufacturer’s instructions.

### In Vivo LPS Challenge

Wild-type C57BL/6 mice were pretreated with NU9056 or saline 0.5 hr earlier, then injected intraperitoneally with LPS (20 mg/kg body weight), after 8 hr, the mice were killed, and the serum concentrations of IL-1β, TNF-α and IL-6 were measured by ELISA.

### MSU-induced peritonitis in vivo

Wild-type C57BL/6 mice were pretreated with NU9056 or saline 0.5 hr earlier, then injected intraperitoneally with 1 mg MSU (dissolved in 500 μL PBS) for 6 hr. Peritoneal cavities were washed with 10 mL ice-cold PBS. The peritoneal lavage fluids were collected and concentrated for ELISA analysis with Amicon Ultra 10 K filter (UFC900308) from Millipore. Peritoneal exudate cells were collected and analyzed by FACS.

### Statistical analysis

The results are expressed as the mean ± SD. Statistical analysis was performed using Student’s t test for two groups with GraphPad Prism, Differences were considered significant when *P<0.05, **P<0.01 and ***P<0.001.

---

**Figure S1.**
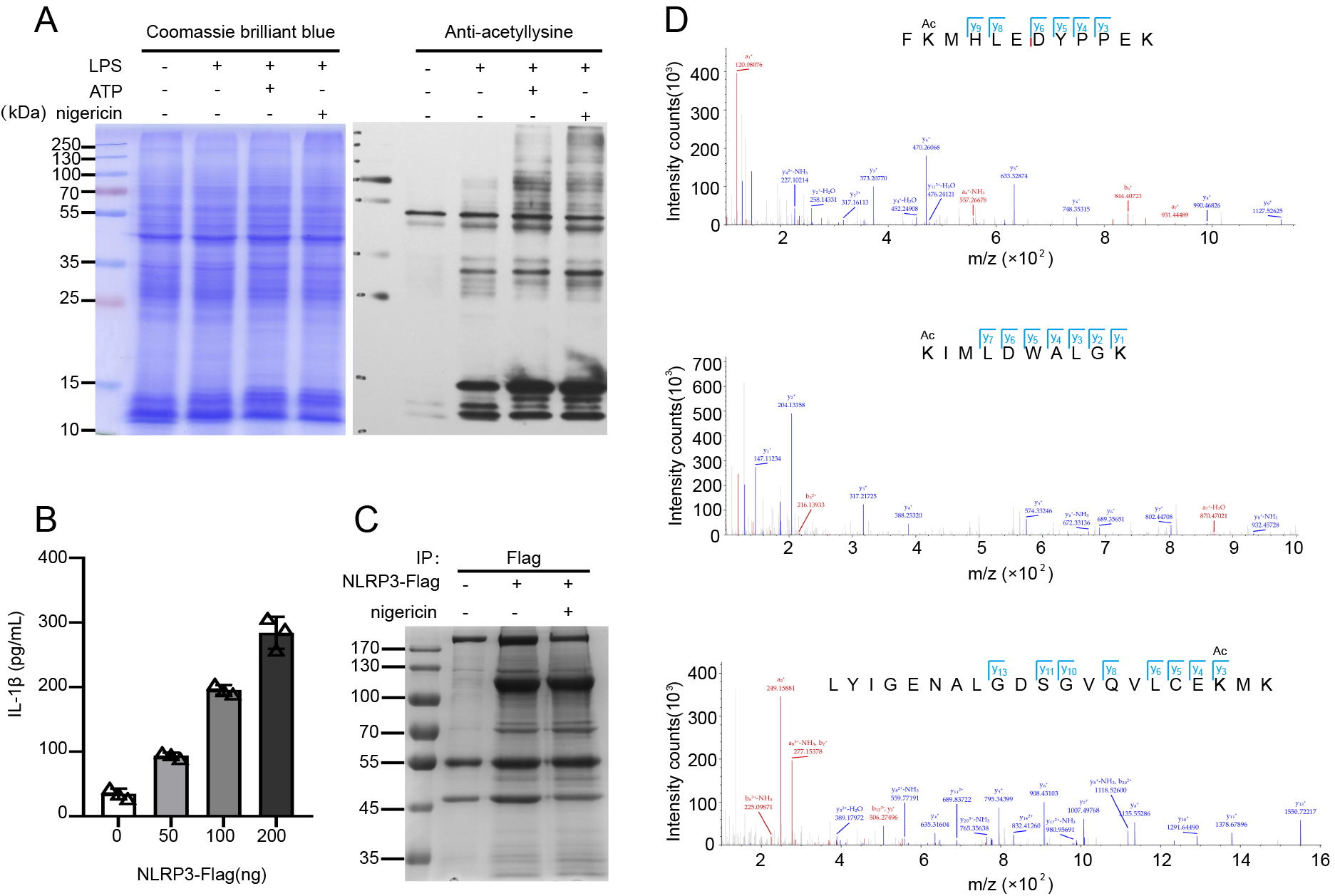
Identification of NLRP3 acetylation. (A) Immunoblot analysis of total acetylation level from lysates of peritoneal macrophages treated with indicated stimuli (B) ELISA of IL-1β in supernatants of HEK293T cells reconstituted with NLRP3 inflammasome and stimulated with nigericin. (C) SDS-PAGE of purified proteins immunoprecipitated by anti-Flag antibody from HEK293T cells. (D) NLRP3 acetylation sites identified with mass spectrometry.

**Figure S2.**
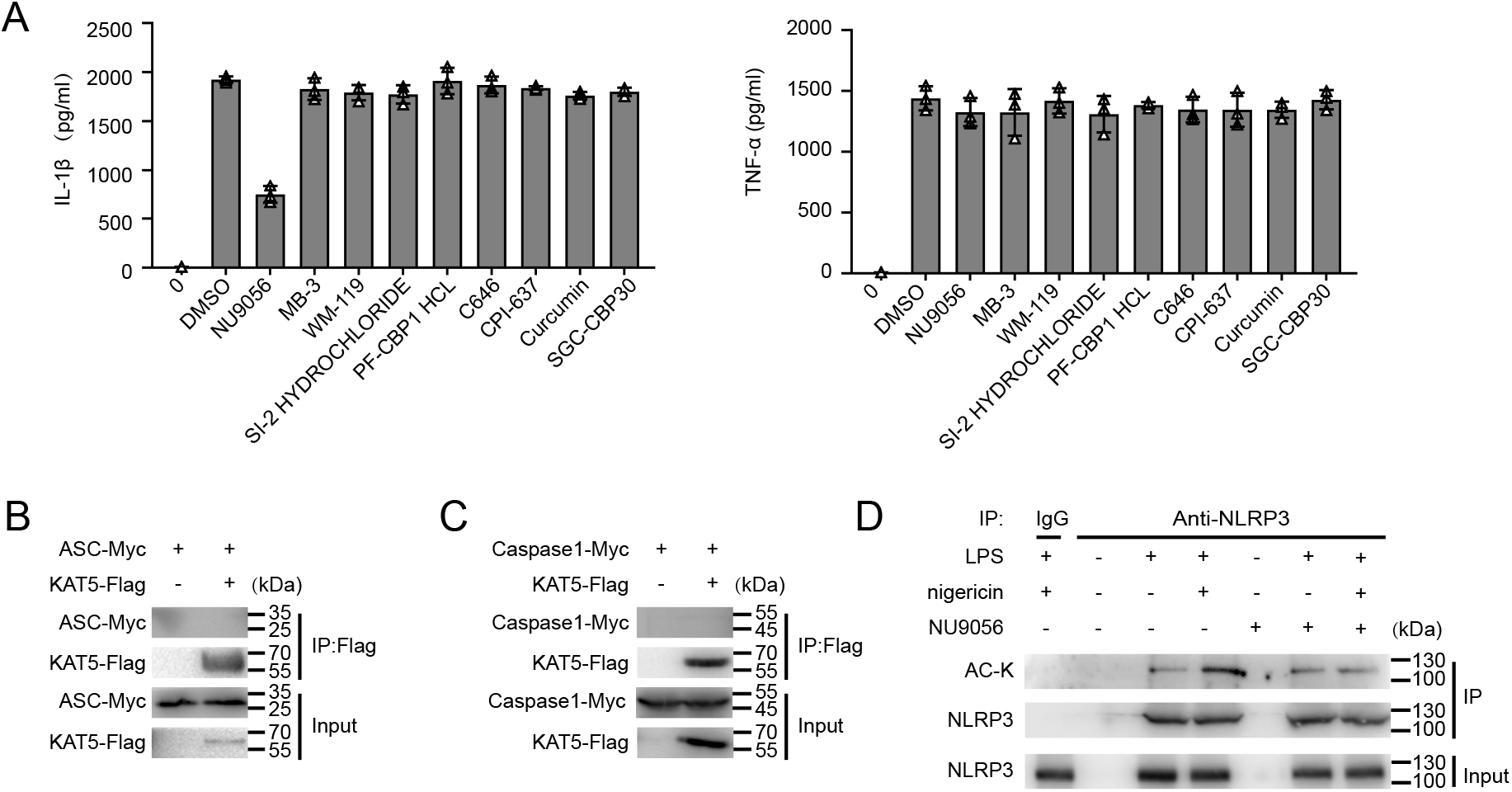
KAT5 is involved in NLRP3 inflammasome activation. (A) Effects of acetyltransferase inhibitors on NLRP3 inflammasome activation. ELISA of IL-1β and TNF-α in supernatants from LPS-primed mouse peritoneal macrophages treated with 1μn various acetyltransferase inhibitors and then stimulated with nigericin. (B and C) Immunoblot analysis of ASC (B) or caspase-1(C) in HEK293T cells transfected to overexpress Myc-tagged ASC(B), caspase-1(C) alone or with Flag-tagged KAT5, assessed before (Input) or after (IP) immunoprecipitation with antibody to Flag. (D) Immunoblot analysis of acetylation level of NLRP3 immunoprecipitated from total lysis of peritoneal macrophage treated with indicated stimuli with or without NU9056. Experiments were repeated at least three times, and representative data are shown. Values, mean±SD.

**Figure S3.**
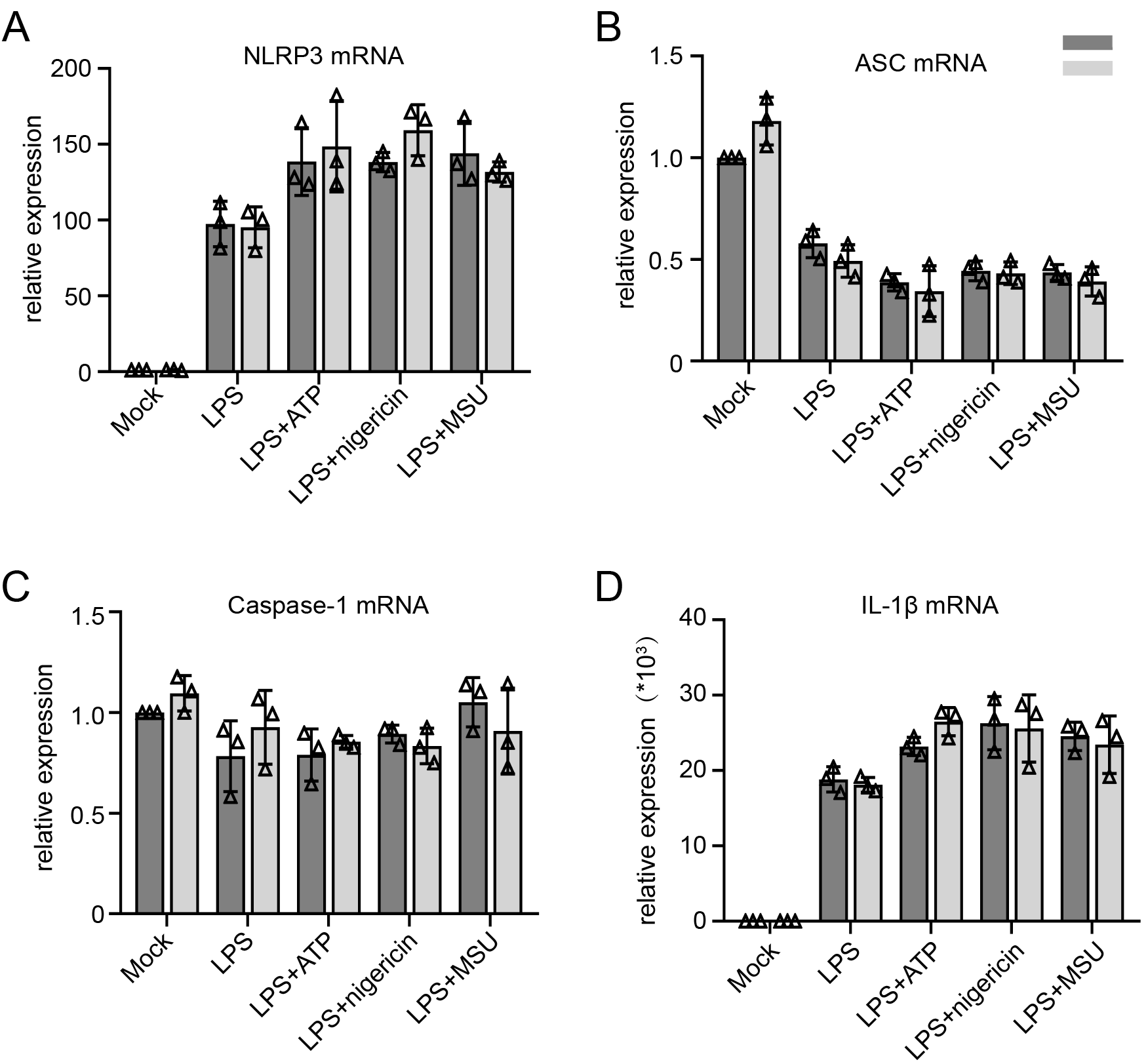
KAT5 barely affects the mRNA expression of NLRP3 inflammasome components. (A-D) Relative NLRP3, ASC, C1, IL-1β mRNA expression of peritoneal macrophage treated with indicated stimuli after transfected with NC-siRNA and KAT5-siRNA. Experiments were repeated at least three times, and representative data are shown. Values, mean± SD.

**Figure S4.**
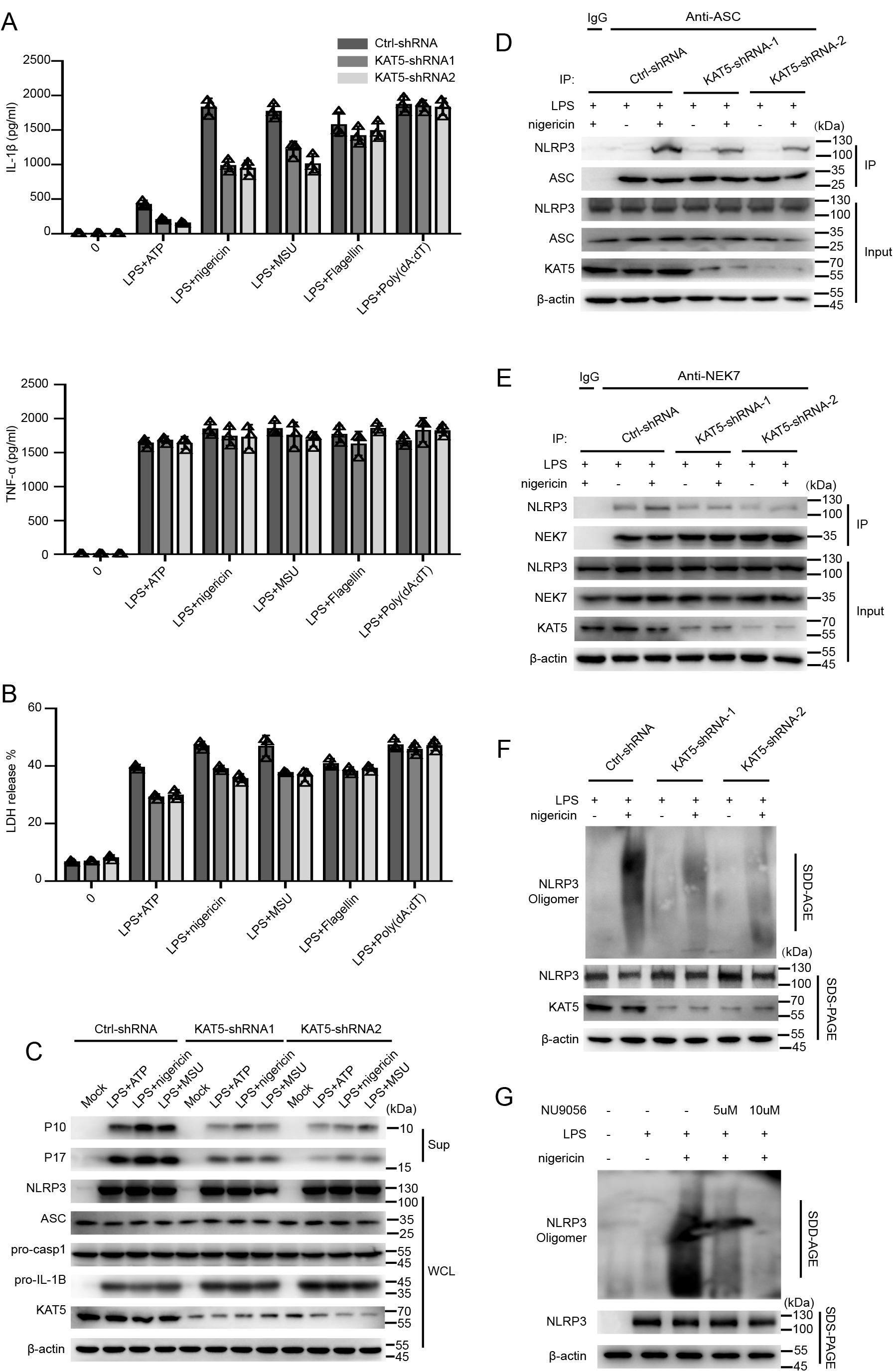
KAT5 promotes NLRP3 inflammasome activation in iBMDMs. (A-C) ELISA of IL-1β and TNF-α(A), release of LDH(B) in supernatants and immunoblot analysis(C) of extracts from LPS primed iBMDMs stably expressing shRNAs targeting KAT5(shRNA-1and shRNA-2), treated with indicated stimuli. (D and E) LPS-primed iBMDMs stably expressing shRNAs targeting KAT5(shRNA-1and shRNA-2), treated with indicated stimuli, immunoprecipitation was performed to detect the interaction between NLRP3 and ASC(D) or NEK7(E). (F) Immunoblot analysis of NLRP3 oligomerization by SDD-AGE or SDS-PAGE assay in LPS-primed iBMDMs stably expressing shRNAs targeting KAT5(shRNA-1and shRNA-2), treated with indicated stimuli. (G) Immunoblot analysis of NLRP3 oligomerization by SDD-AGE or SDS-PAGE assay in peritoneal macrophages treated with various doses of NU9056 and then stimulated with nigericin. Experiments were repeated at least three times, and representative data are shown. Values, mean±SD.

**Figure S5.**
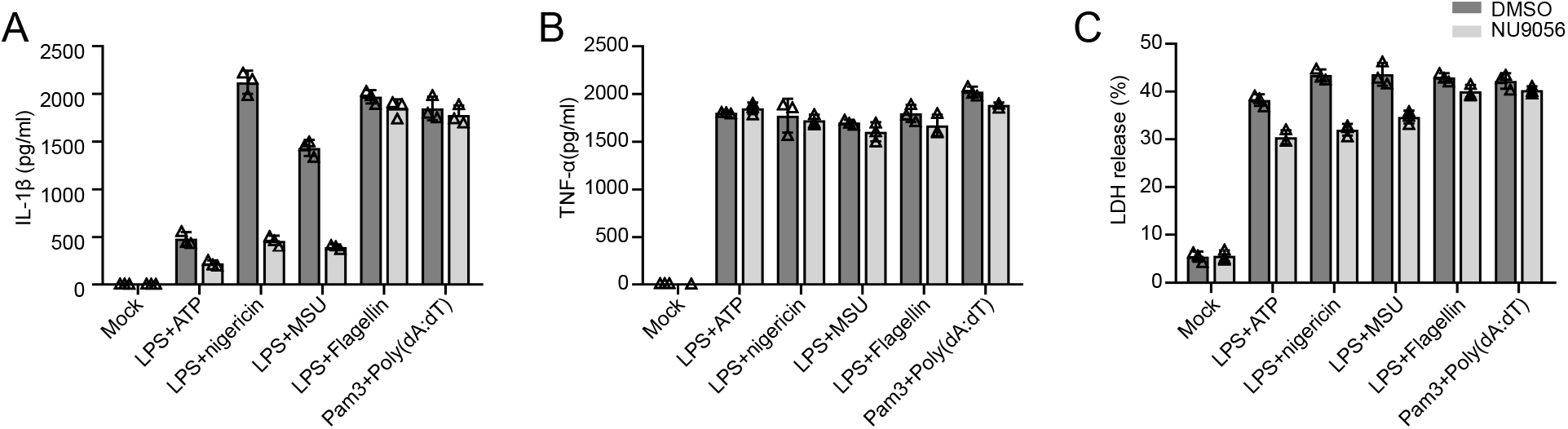
NU9056 specifically blocks NLRP3 inflammasome activation. ELISA of IL-1β (A), TNF-α (B) and release of LDH(C) in supernatants of peritoneal macrophages treated with indicated stimuli with or without NU9056. Experiments were repeated at least three times, and representative data are shown. Values, mean± SD.

