## Supplementary Table1 for "Acetylation is required for NLRP3 self-aggregation and full activation of the inflammasome"

| REAGENT or RESOURCE | SOURCE | IDENTIFIER |
|---|---|---|
| Antibodies | | |
| Anti-Caspase-1 antibody | abcam | ab179515 |
| Anti-IL-1β antibody | RD systems | AF-401-NA; RRID: AB_416684 |
| Anti-KAT5 antibody | abcam | ab23886 |
| Anti-NLRP3 antibody | Adipogen | Cryo-2 |
| Anti-ASC antibody | Adipogen | AL177 |
| Anti-β-actin antibody | Cell Signaling Technology | BH10D10 |
| Anti-NEK7 antibody | abcam | ab133514 |
| Anti-acetyllysine mouse mAb(clone Kac-01) antibody | PTM BIO | PTM-101 |
| Anti-DDDDK-tag | MBL | M185-3L |
| Anti-Myc-tag | MBL | M047-3 |
| Mouse anti GST-Tag mAb | ABclone | AE001 |
| DyLight 488-labeled secondary antibody | InvivoGen | A120-100D2 |
| Alexa Fluor 594-conjugated secondary antibody | InvivoGen | 405326 |
| Chemicals, Peptides, and Recombinant Proteins | | |
| Ultrapure LPS (E. coli 0111:B4) | InvivoGen | tlrl-3pelps |
| Standard LPS (E. coli 0111:B4) | InvivoGen | tlrl-eblps |
| Pam3CSK4 | InvivoGen | tlrl-pms |
| ATP | InvivoGen | tlrl-atpl |
| Nigericin | InvivoGen | tlrl-nig |
| MSU | InvivoGen | tlrl-msu |
| FLA-ST | InvivoGen | tlrl-stfla |
| Poly(dA:dT) naked | InvivoGen | tlrl-patn |
| Imiquimod | Invivogen | R837 |
| Lipofectamine 3000 Transfection Reagent | ThermoFisher Scientific | L3000015 |
| NU9056 | Tocris | 4903 |
| Cell Lysis Buffer | Cell Signaling Technology | 9803 |
| Mouse immunoglobin IgG protein | Abcam | ab198772 |
| Protein A/G PLUS-Agarose | Santa cruz | sc-2003 |
| Glutathione Sepharose™ 4B | GE Healthcare | 17-0756-01 |
| Pierce™ Anti-c-Myc Agarose | ThermoFisher Scientific | 20168 |
| Anti-Flag affinity gel | Sigma | A2220 |
| pLenti-CRISPR v2 | Addgene | #52961 |
| First-Strand cDNA Synthesis SuperMix | TransGen Biotech | AT34 |
| SYBR qPCR Master Mix | Vazyme Biotech | Q711-02/03 |
| Critical Commercial Assays | | |
| Mouse IL-1β ELISA kit | eBioscience | 88-7013 |
| Mouse TNF-α ELISA kit | eBioscience | 88-7324 |
| LDH Cytotoxicity Assay Kit | Beyotime | C0017 |

| | | |
|---|---|---|
| **Experimental Models: Cell Lines** | | |
| Mouse Macrophages | Prepared in B.L. Lab | Described in current manuscript |
| HEK293T cells | American Type Culture Collection (Manassas, VA) | N/A |
| NLRP3-/- iBMDM cells | Prepared in B.L. Lab | Described in current manuscript |
| **Experimental Models: Organisms/Strains** | | |
| C57BL/6 mice | Hunan SJA Laboratory Animal Co.Ltd | N/A |
| **Oligonucleotides** | | |
| CCACACUGCAGUAUCUCAATT | Sangon Biotech Co. | KAT5-specific siRNA |
| UUCUCCGAACGUGUCACGUTT | Sangon Biotech Co. | Control siRNA |
| 5'-CTGCAACGCCACTTGACCAAA-3' | Genechem Co | KAT5-specific shRNA1 |
| 5'- CTGCTTATTGAGTTCAGCTAT -3' | Genechem Co | KAT5-specific shRNA2 |
| 5'-TTCTCCGAACGTGTCACGT-3' | Genechem Co | Control shRNA |
| Fwd：5'-TCCCGGTCCAGATCACACTC-3'<br>Rev：5'-ACCTTCCGTTTCGTTGAGCG-3' | Sangon Biotech Co. | KAT5-specific primers |
| Fwd：5'-CTTGTCAGGGGATGAACTCAAAATT-3'<br>Rev：5'-GCCATACGACTCCAG ATAGTAGC-3' | Sangon Biotech Co. | ASC-specific primers |
| Fwd：5'-ACAAGGCACGGG ACCTATG-3'<br>Rev：5'-TCCCAGTCAGTCCTGGAAATG-3' | Sangon Biotech Co. | Caspase-1-specific primers |
| Fwd：5'-TGGATGGGTTTGCTGGGAT-3'<br>Rev：5'-CTGCGTGTAGCGACTGTTGAG-3' | Sangon Biotech Co. | NLRP3-specific primers |
| Fwd：5'- GCAACTGTTCCTGAACTCAACT-3'<br>Rev：5'- ATCTTTTGGGGTCCGTCAACT-3' | Sangon Biotech Co. | IL-1β-specific primers |
| Fwd：5'-AGTGTGACGTTGACATCCGT-3'<br>Rev：5'-GCAGCTCAGTAACAGTCCGC-3' | Sangon Biotech Co. | β-actin-specific primers |
| **Software and Algorithms** | | |
| Graphpad Prism 8 software | Graphpad Prism 8 software | N/A |
| Adobe Illustrator CS6 | Adobe | N/A |
| Adobe Photoshop CC | Adobe | N/A |
| Microsoft Excel | Microsoft | N/A |
